# Evidence for evolutionary adaptation of mixotrophic nanoflagellates to warmer temperatures

**DOI:** 10.1101/2022.02.03.479051

**Authors:** Michelle Lepori-Bui, Christopher Paight, Ean Eberhard, Conner M. Mertz, Holly V. Moeller

## Abstract

Mixotrophs, organisms that combine photosynthesis and heterotrophy to gain energy, play an important role in global biogeochemical cycles. Metabolic theory predicts that mixotrophs will become more heterotrophic with rising temperatures, potentially creating a positive feedback loop that accelerates carbon dioxide accumulation in the atmosphere. Studies testing this theory have focused on phenotypically plastic (short-term) thermal responses of mixotrophs. However, as small organisms with short generation times and large population sizes, mixotrophs may rapidly evolve in response to climate change. Here we present data from a 3-year experiment quantifying the evolutionary response of two mixotrophic nanoflagellates to temperature. We found evidence for adaptive evolution through increasing growth rates in the obligately mixotrophic strain, but not in the facultative mixotroph. All lineages showed trends of increased carbon use efficiency, flattening of thermal reaction norms, and a return to homeostatic gene expression. Generally, mixotrophs evolved reduced photosynthesis and higher grazing with increased temperatures, suggesting that evolution may act to exacerbate mixotrophs’ effects on global carbon cycling.

## 1. Introduction

Mixotrophs, organisms that use a combination of autotrophy and heterotrophy to gain energy and nutrients, are increasingly recognized as omnipresent members of planktonic food webs and regulators of global biogeochemical cycles (Mitra *et al*. 2014; Worden *et al*. 2015; Caron 2016; Ward & Follows 2016). “Constitutive mixotrophs” are chloroplast-bearing protists that have retained the ability to eat (Stoecker 1998; Mitra *et al*. 2016). These mixotrophs occur on a spectrum of metabolic strategies, ranging from primarily phototrophic (feeding when nutrients needed for photosynthesis are limiting) to primarily heterotrophic (photosynthesizing when prey are limiting) (Stoecker 1998). Though different species of mixotrophs may favor one mode of carbon acquisition over the other in ideal conditions, the balance between autotrophy and heterotrophy is also affected by environmental factors including temperature, light, and prey availability.

As a result of their flexible metabolism, mixotrophs may act as either carbon sources or sinks. For example, mixotrophs contribute substantially to primary production in mature ecosystems (Mitra *et al*. 2014) where they can account for over 80% of total chlorophyll (Zubkov & Tarran 2008). They can also drive carbon remineralization as the dominant grazers in oligotrophic gyres (Hartmann *et al*. 2012). Ocean ecosystem models predict that incorporating mixotrophy can promote the accumulation of biomass in larger size classes, increasing estimates of carbon export via the biological carbon pump by 60% (Ward & Follows 2016). However, accurate model predictions require a better understanding of mixotroph metabolic flexibility, particularly in the face of ocean warming.

Rising ocean temperatures due to climate change will fundamentally affect oceanic ecosystems by altering the metabolic functions of marine organisms (Gillooly *et al*. 2001). According to the metabolic theory of ecology, metabolic rates increase exponentially with temperature (Brown *et al*. 2004). Because heterotrophic processes are more sensitive to temperature increases than photosynthetic processes (Allen *et al*. 2005; Rose & Caron 2007), mixotrophs are predicted to become more heterotrophic at higher temperatures (Allen *et al*. 2005; Wilken *et al*. 2013), increasing their emission of carbon dioxide. Further, mixotrophs are dominant in the low-nutrient stratified water of subtropical gyres (Hartmann *et al*. 2012; Mitra *et al*. 2016), which are expected to expand with climate change (Polovina *et al*. 2008). Rising temperatures are also associated with decreases in body size (Gillooly *et al*. 2001; Malerba & Marshall 2020), which reduces sinking rates, leading to cascading effects to carbon export by the biological carbon pump. Together, these changes could increase carbon dioxide accumulation in the atmosphere, generating a positive feedback loop.

Relatively few studies have tested the hypothesis that mixotrophs will become more heterotrophic with increased temperatures. Wilken *et al*. (2013) found a shift towards heterotrophy at higher temperatures in a primarily phagotrophic freshwater mixotroph, *Ochromonas sp*., after 2-4 weeks of acclimation to new temperatures. Conversely, Princiotta *et al*. (2016) found the opposite effect after 5 days of thermal acclimation in an obligately phototrophic freshwater mixotroph, *Dinobryon sociale*. These contrasting results show that predicting mixotrophs’ response to increased temperature is complicated by many factors, including the diversity of mixotrophic organisms and how they balance their metabolism. Furthermore, due to the short time scale of these experiments (between 5 days and 4 weeks for slower-growing organisms), these results represent the organism’s phenotypic plasticity, or the ability of a single genotype to exhibit different traits as a function of abiotic conditions. However, due to their short generation times, fast growth rates, and large population sizes, microbes may rapidly adapt (via evolutionary changes in genotype) to changing conditions. For example, a growing body of evolutionary experiments has shown that some phytoplankton are capable of adaptive evolution within several hundred generations (Listmann *et al*. 2016; Padfield *et al*. 2016; O’Donnell *et al*. 2018; Schaum *et al*. 2018; Barton *et al*. 2020). In some cases, these adaptations reversed short-term responses to increasing temperature, such as through the reduction of respiratory costs (Padfield *et al*. 2016; Barton *et al*. 2020). Although high taxonomic diversity mean lineages likely vary in their responses (Collins *et al*. 2014), to our knowledge, no mixotrophs have been similarly experimentally evolved.

Here, we quantified the evolutionary responses of mixotrophs to temperature change and related these responses to mixotrophic contributions to carbon cycling in the oceans. Specifically, we asked: Do mixotrophs adapt to different temperatures? What changes can be observed in carbon cycle-relevant traits when comparing their plastic and evolved responses? And what are some of the potential mechanisms for adaptation? We experimentally evolved two related strains of mixotrophic nanoflagellates—one obligate mixotroph, requiring both light and prey, and one facultative phototroph, requiring prey but able to grow in darkness—to different temperature treatments for three years, to quantify adaptation. We varied light availability to manipulate selection for photosynthesis, and measured carbon cycle-relevant traits including metabolic rates (photosynthesis, grazing, and respiration), photosynthetic parameters, and cell size. We found evidence for adaptive evolution to both hot and cold temperatures in the obligately mixotrophic strain, but only under high light conditions. Although differences in fitness over time were more variable in the facultative mixotroph, evolution increased carbon use efficiency and reversed some of the short-term stress responses of control lineages. All lineages showed evolved responses in carbon cycle-relevant traits at the end of our experiment that could exacerbate mixotroph contributions to climate change.

## 2. Materials and Methods

### Mixotroph cultures and maintenance

We experimentally evolved two marine lineages from the genus *Ochromonas*, a widely distributed group of mixotrophic nanoflagellates. These cultures, purchased from the National Center for Marine Algae and Microbiota (NCMA, Bigelow Laboratory, East Boothbay, ME), represent different degrees of metabolic flexibility: Strain CCMP 1393 is obligately mixotrophic (requiring both light and prey to survive), and Strain CCMP 2951 is facultatively phototrophic (requiring prey but able to grow in darkness) (Wilken *et al*. 2020). Cultures were maintained in K medium (Keller *et al*. 1987) made by adding pre-mixed nutrients (NCMA) to 0.2 μm filtered coastal seawater. Stock cultures were kept at the ancestral temperature of 24°C, with a 12:12 light:dark cycle and acclimated to the two experimental light levels, 100 and 50 μmol quanta·m^−2^·s^−1^, for at least 5 weeks prior to the start of the experiment.

### Evolution experiment

We conducted a long-term evolution experiment testing mixotroph responses to both lower (18°C) and higher (30°C) temperatures. In March 2018, we initiated six evolutionary replicates for each temperature treatment with a sub-population of 10,000 cells from stock cultures maintained at 24°C (hereafter, “control”) at each light level (Fig. 1A). This initial population size was chosen over using a clonal isolate to avoid genetic bottlenecks, increase the probability of favorable mutations, and support the long-term stability of the cultures (Wahl *et al*. 2002; Elena & Lenski 2003; Malerba & Marshall 2020). We monitored cell density weekly by counting a live sub-sample of each lineage using a Guava easyCyte flow cytometer (Luminex Corporation, Austin, USA), distinguishing *Ochromonas* cells using forward scatter (a proxy for cell size) and red fluorescence (a measure of photosynthetic pigmentation). Evolving cultures were transferred in a 1:15 mL dilution of fresh media in 25 mL Culture Flasks (Genesee Scientific, Part No. 25-212, San Diego, CA, USA) every subsequent 2-4 weeks (depending on population density) to maintain exponential growth and maximize adaptive potential (Elena & Lenski 2003).

**Figure 1.**
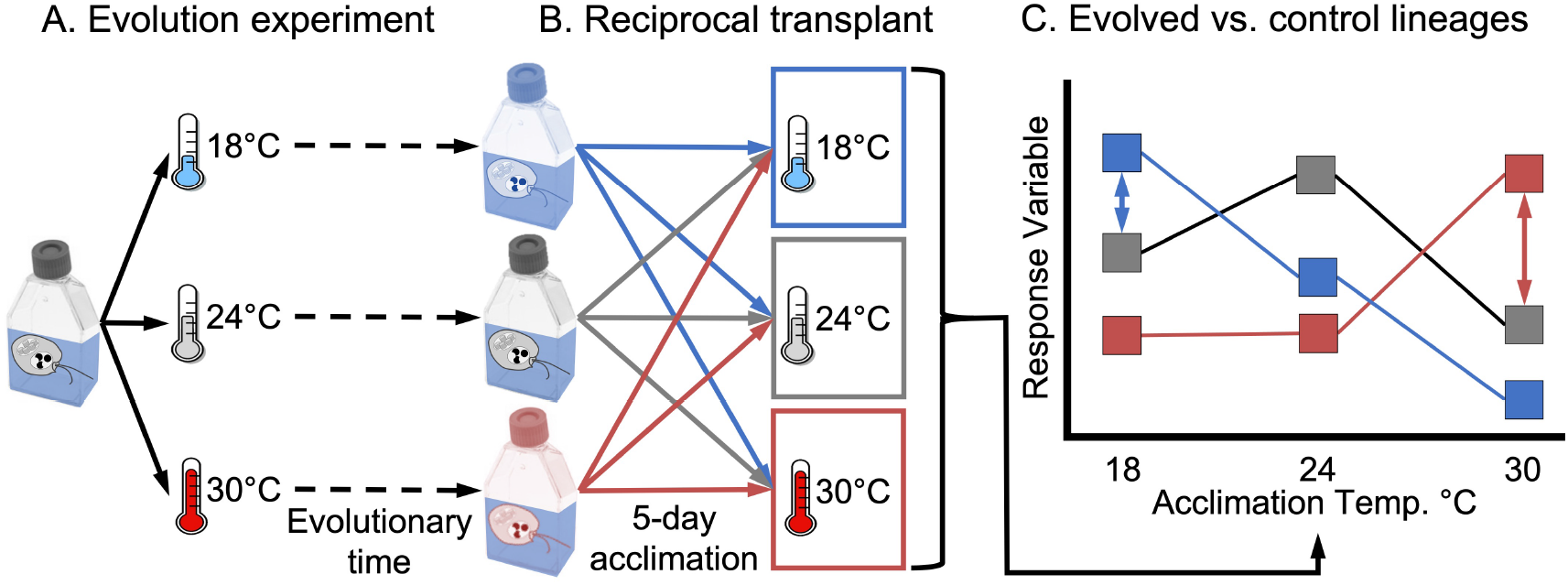
Conceptual diagram of experimental design. A. Genetically similar ancestral cultures were split and exposed to three treatment temperatures (18°C, 24°C (control), and 30°C) for evolutionary time scales. B. Reciprocal transplant assays were performed periodically, wherein aliquots of each strain were transplanted to each treatment temperature for an acclimation period of 5 days prior to trait tests to differentiate between adaptation and plasticity. C. Hypothetical data points demonstrate the thermal reaction norms for each treatment lineage that result from reciprocal transplant assays. In particular, we can compare the cold-evolved (blue squares) and hot-evolved (red squares) to the control lineages (gray squares). Differences between these (blue and red double-sided arrows) shows evolutionary response, while no differences in thermal reaction norms (not pictured) would indicate plasticity.

### Reciprocal transplant assays

To differentiate between evolution and plasticity, we periodically transplanted evolving lineages to all treatment temperatures and quantified metabolic traits relevant to the carbon cycle (Fig. 1B-C). Aliquots of evolving cultures were transferred to new temperatures for a 5-day acclimation to overcome transfer shock before experimental measurements began. We measured growth rate every three months to test overall fitness; photosynthetic, grazing, and respiration rates every six months to quantify carbon budgets; and cellular carbon and nitrogen content (yearly after Year 1 of the experiment) to determine cell size and stoichiometry. All physiological measurements were made on cells in exponential growth phase and between 5-10 days of temperature acclimation. All analyses were performed using the software package R (R Core Team 2020). Data and code are available at: https://github.com/mleporibui/OchEvo. [Note to reviewers: A permanent DOI will be created when the manuscript is accepted, and code is finalized.]

### Growth rate and generation time

Growth assays were conducted in 24-well plates (VWR, Part No. 10062-896, Radnor, PA, USA). On Day 6 of temperature acclimation, *Ochromonas* lineages were inoculated into 2.5 mL of media at an initial density of 20,000 cells·mL^−1^ and counted daily for 4 days using a flow cytometer. Growth rates were calculated by fitting a linear model to the natural log of population size (R function *lm*). Generation time for experimental (18°C and 30°C) lineages was calculated over the course of the experiment using growth rates specific to each strain, temperature, and time step. We divided log(2) by the growth rate (interpolated over time using R function *smooth.spline*) to obtain doubling time, then integrated over time to calculate generations elapsed for each lineage between each growth rate measurement.

### Cellular C content

Cellular carbon (C) and nitrogen (N) content were measured using an elemental analyzer (Model CEC 440HA, Exeter Analytical, Coventry, UK). Known volumes and densities of *Ochromonas* cultures were filtered onto pre-combusted GF/F filters (Whatman Part No. 1825-025 Whatman Cytiva, Marlborough, MA, USA), acidified to remove inorganic carbonates, and dried before combustion. To control for the biomass of coexisting bacteria, bacteria-only cultures (made by inoculating media with bacteria isolated from stock *Ochromonas* cultures) at each temperature were filtered, acidified, dried, and combusted. Bacteria were enumerated by plating on Difco™ Marine Broth 2216 (Becton, Dickinson and Company, Sparks MD 21152 USA) agar (VWR, Part No. J637, Radnor, PA, USA) plates. Colony forming units were counted after 7 days incubated at 24°C, and average bacterial C content was calculated. In *Ochromonas* cultures, bacteria were similarly enumerated, and bacterial contribution of C was subtracted from the mixed-culture measurements prior to calculating *Ochromonas* cellular C content.

### Photosynthesis and respiration

Photosynthesis and respiration rates were measured using oxygen sensor spots (PyroScience, Aachen, Germany) and FireStingO_2_ optical oxygen meters (Pyroscience, Jallet *et al*. 2016). *Ochromonas* were sealed into airtight glass vials with sensor spots and magnetic stir bars to keep cultures well-mixed. We monitored oxygen levels within vials continuously for 3 hours in light and 2 hours in darkness. To subtract respiratory contributions from coexisting bacteria, we measured respiration rates of bacteria-only cultures (see cellular C content methods) at all treatment temperatures. Temperature-specific bacterial respiration rates were removed from mixed cultures before calculating *Ochromonas* rates. Net photosynthesis and dark respiration rates were calculated by fitting a linear model to change in oxygen (O_2_) measurements (R function *lm*) in light and darkness, respectively. Gross photosynthetic rates were computed as the sum of net photosynthesis and dark respiration, assuming that respiration rates did not change with light. We used the equation from Barton *et al*. (2020) to convert metabolic rates (*b*) from units of O_2_ to μgC (Equation 1).

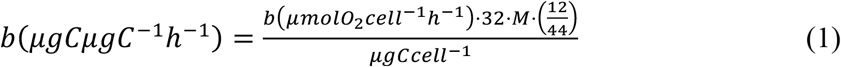

Equation 1 uses the molecular weights of O_2_, C, and carbon dioxide (CO_2_), and the species-specific assimilation quotient (M) from Falkowski *et al*. (1985). M describes the conversion between C and O_2_ through consumption or fixation withing a cell using its C:N ratio.

### Grazing

We measured *Ochromonas* grazing rates by offering mixotrophs heat-killed fluorescently labeled bacteria (FLB; *Escherichia coli* – K-12 strain-Bioparticles®, Alexa Fluor®488 conjugate, Molecular Probes, Invitrogen, Waltham, MA, USA) as prey. To construct grazing functional response curves, we inoculated FLB at a range of concentrations between 0 and 4 million cells·mL^−1^ into *Ochromonas* cultures, as well as into sterile media as controls. After 1 h of grazing, final concentrations of *Ochromonas* and FLB were enumerated using forward scatter, red-fluorescence, and yellow-fluorescence measurements on a flow cytometer. Grazing rates were calculated using methods of Jeong and Latz (1994) for each FLB concentration. We fit both Holling Type I (linear) and Type II (saturating) functional response curves to the data and used Akaike Information Criterion (AIC) values to determine which response fit best. Grazing rates were calculated at average prey densities, which were determined as average bacteria per treatment, enumerated as described in cellular C content methods.

### Photosynthetic traits

We measured electron transport rate (ETR) and photosynthetic efficiency (Fv/Fm) using a mini–Fluorescence Induction and Relaxation (miniFIRe) system (custom built by M. Gorbunov, Rutgers University, New Brunswick, NJ, USA). We quantified photosynthetic rate as a function of irradiance according to Gorbunov *et al*. (1999). Photosynthetic efficiency was measured as the ratio of variable to maximum fluorescence. ETR was measured at light intervals between 0 and 1000 μmol quanta·m^−2^·s^−1^ to generate photosynthesis-irradiance curves, which were fit with the photosynthesis-irradiance equation of Jassby and Platt (1976) (Equation 2) using non-linear least squares regression (R function *nls*).

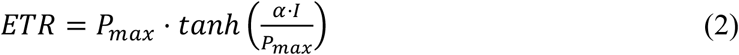

In Equation 2, ETR is calculated using maximum ETR (*P_max_*), the initial slope of ETR to light (α), and the incident irradiance (*I*). We extracted chlorophyll-*a* (chl-*a*) by incubating a known number of cells captured on a GF/F filter (Whatman Part No. 1825-025, Whatman Cytiva, Marlborough, MA, USA) overnight in 90% acetone at 4°C, then quantified it using a Trilogy fluorometer with a 460nm LED (Turner Designs, San Jose, CA, USA). We used a linear model to calibrate between flow cytometry red fluorescence to extracted chl-*a* content (Fig. S1).

### Thermal reaction norms

To obtain a more complete picture of the thermal sensitivity of metabolic traits in evolved lineages, we measured thermal reaction norms (TRNs) of photosynthesis and bacterivory. We performed photosynthesis-irradiance curves or grazing assays as described above at a range of temperatures between 3°C to 44°C. These thermal assays represent short-term responses of cells to temperature where samples were incubated at the assay temperature only for the duration of the assay (15 minutes of dark acclimation for photosynthesis-irradiance curves; 60 minutes of incubation with FLB for grazing assays). For photosynthesis-irradiance curves, to rapidly bring cells to their incubation temperature, we diluted 1 mL of *Ochromonas* culture with 4 mL of sterile, filtered seawater at the assay temperature. We only collected TRN data from the obligate mixotroph (Strain 1393) evolved at high light because this strain showed the strongest adaptive response to temperature. Thermal reaction norms for ETR were fit using a reparameterized version of the Norberg-Eppley equation (Equation 3) from the R package “growthTools” (Kremer, 2021).

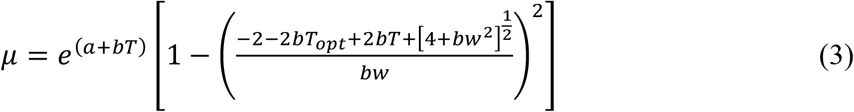

In Equation 3, *μ* is the metabolic trait, *T* is temperature, *T_opt_* is the optimum temperature, *a* affects the y-intercept, *b* affects thermal scaling, and *w* describes the thermal niche width. For photosynthetic efficiency and grazing, we added smoothed conditional means (R function *ggplot2::geom_smooth*).

### Transcriptome sequencing

Finally, we measured gene expression in the obligate mixotroph (Strain 1393) evolved at high light to better understand cellular mechanisms underlying observed adaptive responses. To contrast evolved and acclimatory responses, we collected transcriptomes from lineages evolving at all temperatures, and from control lineages evolving at 24°C but acclimated to 18°C or 30°C. In week 141 of the experiment, we inoculated exponentially growing *Ochromonas* cells into 130 mL volumes of culture media in 250 mL tissue culture flasks (VWR Part # 10062-860). We incubated these cultures for 48 hours at their evolutionary temperatures (to allow cells to overcome transfer shock), before transplanting acclimation treatments to their new temperatures. Evolving treatments remained at their initial temperatures. After seven days (the mean acclimation time of our reciprocal transplant studies; see above), we filtered the cultures onto 0.8μm pore size polycarbonate filters (Millipore ATTP04700, Millipore Sigma, Darnstadt, Germany), immediately flash-froze samples in liquid nitrogen, and then stored them at −80°C until RNA extraction (within one week). We collected one transcriptome from each evolving or acclimated lineage (= 5 treatments x 6 replicates for a total of 30 samples), except for lineages evolving at 18°C for which we collected technical triplicates (i.e., inoculated 3 replicate 130 mL volumes of media, and collected 3 replicate transcriptomes) to contrast the range of gene expression in technical replicates with that contained amongst biological replicates.

We extracted samples using an RNeasy Plant Mini Kit (Qiagen, Hilden, Germany). Cells were physically disrupted by adding 2.8 mm ceramic beads (Qiagen) and 400 μL Buffer RLT with 10 μL/mL β-mercaptoethanol (Qiagen), then vortexed for 30 seconds. Following cell lysis, RNA extraction proceeded according to manufacturer’s instructions. Samples were sequenced at the UC Davis Genome Center (Davis, CA, USA). We assembled uncorrected reads *de novo* with RNA SPAdes (v3.13.0; default parameters, k = 49, 73) (Bankevich *et al*. 2012) and used TransDecoder (v5.5.0) to translate assemblies into protein sequences. We compared our predicted proteins with the NCBI RefSeq database using Diamond v2.02 (-p 32 −b 8 −c 1) (Buchfink *et al*. 2015); Sequences that were identified as bacterial (>90% sequence identity and >80% query coverage) were considered contaminants and removed from further analysis. We then assessed assembly completeness using Busco v 5.0.0 (Simão *et al*. 2015), and used KEGG GhostKoala (Kanehisa *et al*. 2016) to perform preliminary annotations. Read mappings to nucleotide transcripts were quantified with Salmon 0.12.0 (-l A --validateMappings --gcBias) (Patro *et al*. 2017), and differential expression was analyzed with the R package DEseq2 v1.28.1 (Love *et al*. 2014). Differential expression was calculated with Approximate Posterior Estimation for GLM (apeglm) (Zhu *et al*. 2019).

To quantify the effects of evolutionary history on gene expression, we first confirmed that single replicate transcriptomes were sufficient to capture variation in gene expression within treatment group. We did this by contrasting differential expression within technical replicates and across biological replicates for lineages evolved at 18°C. Next, we identified genes with evidence of differential expression (>2x change in expression; adjusted p-value < 0.1) across any treatment group and asked whether differentially expressed genes tended to be up- or down-regulated in response to temperature. To study genes linked to thermal evolution, we selected genes with differential expression between *either* lineages evolved at 18°C and those acclimated to 18°C *or* lineages evolved at 30°C and those acclimated to 30°C. We then contrasted expression in this subset of genes in thermally evolved or acclimated lineages with control lineages.

## 3. Results

### 3.1 Obligate mixotroph growth rates evolved in response to temperature

We found evidence for thermal adaptation in the obligate mixotroph, *Ochromonas* Strain 1393, when it was evolved in high light (100 μmol quanta·m^−2^·s^−1^) at both cold and hot temperatures (Fig. 2A). Within fifty generations, evolving lineages grew faster than the acclimated controls (lineages evolving at ancestral temperatures and assayed at the evolutionary temperature, Fig. S2). Evidence for adaptation in the obligate mixotroph was weaker at the lower light level (50 μmol quanta·m^−2^·s^−^), which showed significant relative increases in growth rate at cold temperatures after about 50 generations, but mixed evidence for adaptation at hot temperatures (4 out of 11 reciprocal transplant experiments showed increases in growth, Student’s t-test, Fig. 2B). Growth rates of the facultative mixotroph, *Ochromonas* Strain 2951, were much more variable, and did not show consistent evidence of evolution in any direction (Fig. 2C-D).

**Figure 2.**
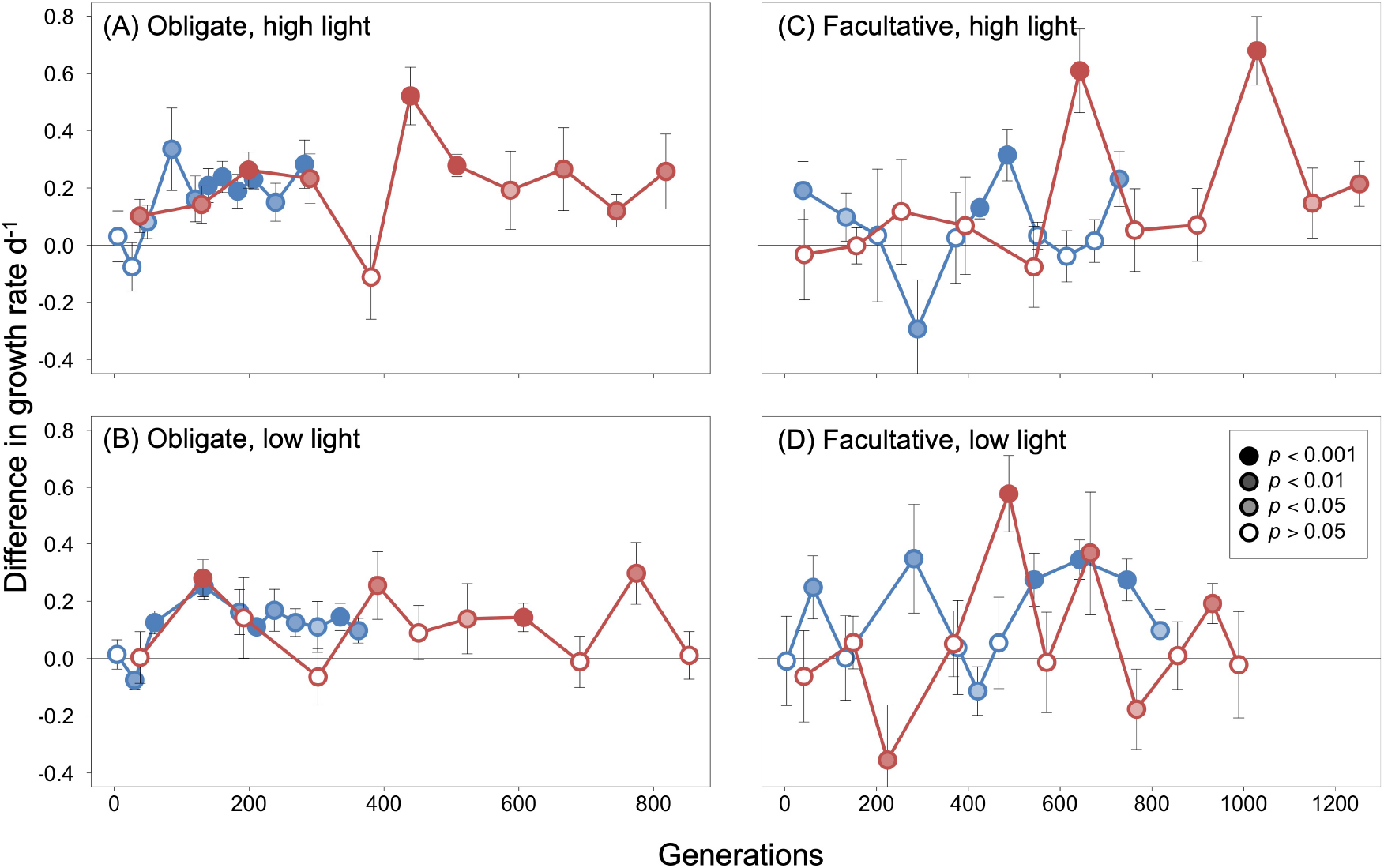
Thermal adaptation in obligate mixotrophs. To test for evolutionary responses, we computed the difference in growth rates between experimental and acclimated control lineages at hot (red) and cold (blue) temperatures. Positive values are evidence of adaptive evolution (experimental lineages growing faster than acclimated control lineages assayed at the same temperature). The x-axes are scaled by growth rate to show time in number of generations the evolving strains experienced at their evolutionary temperatures. Data are shown for both mixotroph strains—the obligate mixotroph Strain 1393 (left column) and the facultative mixotroph Strain 2951 (right column) – and both light levels – high light (100 μmol quanta·m^−2^·s^−1^ top row) and low light (50 μmol quanta·m^−2^·s^−1^, bottom row) for each reciprocal transplant experiment. Point coloration indicates significance ranging from *p*<0.001 (darkest points; one sample Student’s t-test) to *p*>0.05 (white).

### 3.2 Evolution affects mixotroph traits

Although only the obligate mixotroph showed strong evidence for adaptation over time (in the form of increases in growth rates relative to the acclimated controls), all mixotroph strains displayed evolutionary responses to temperature in carbon cycle-relevant traits. Generally, evolutionary thermal responses were less variable than phenotypically plastic ones (Fig. 3). By year three of the experiment, most lineages showed some evidence of adaptation through increased growth rate compared to acclimated controls (Fig. 3, top row), though note that this represents 3 time points that are part of a more equivocal trend in the facultative mixotroph. As a result, evolution produced a general trend of increasing growth rates with ambient temperature (true after evolution in all lineages except the facultative mixotroph at low light, Fig. 3, top row). For obligate mixotrophs, cellular C content (a proxy for cell size) generally decreased with temperature in evolved lineages (Fig. 3, second row), though this was driven by larger cell sizes at the coldest temperature. In all cases, mixotrophs evolved smaller cell sizes at the hottest temperatures relative to phenotypic plasticity in the control (Fig. 3, second row, compare red and gray points), but these cell sizes were not always smaller than those evolved at cooler temperatures.

**Figure 3.**
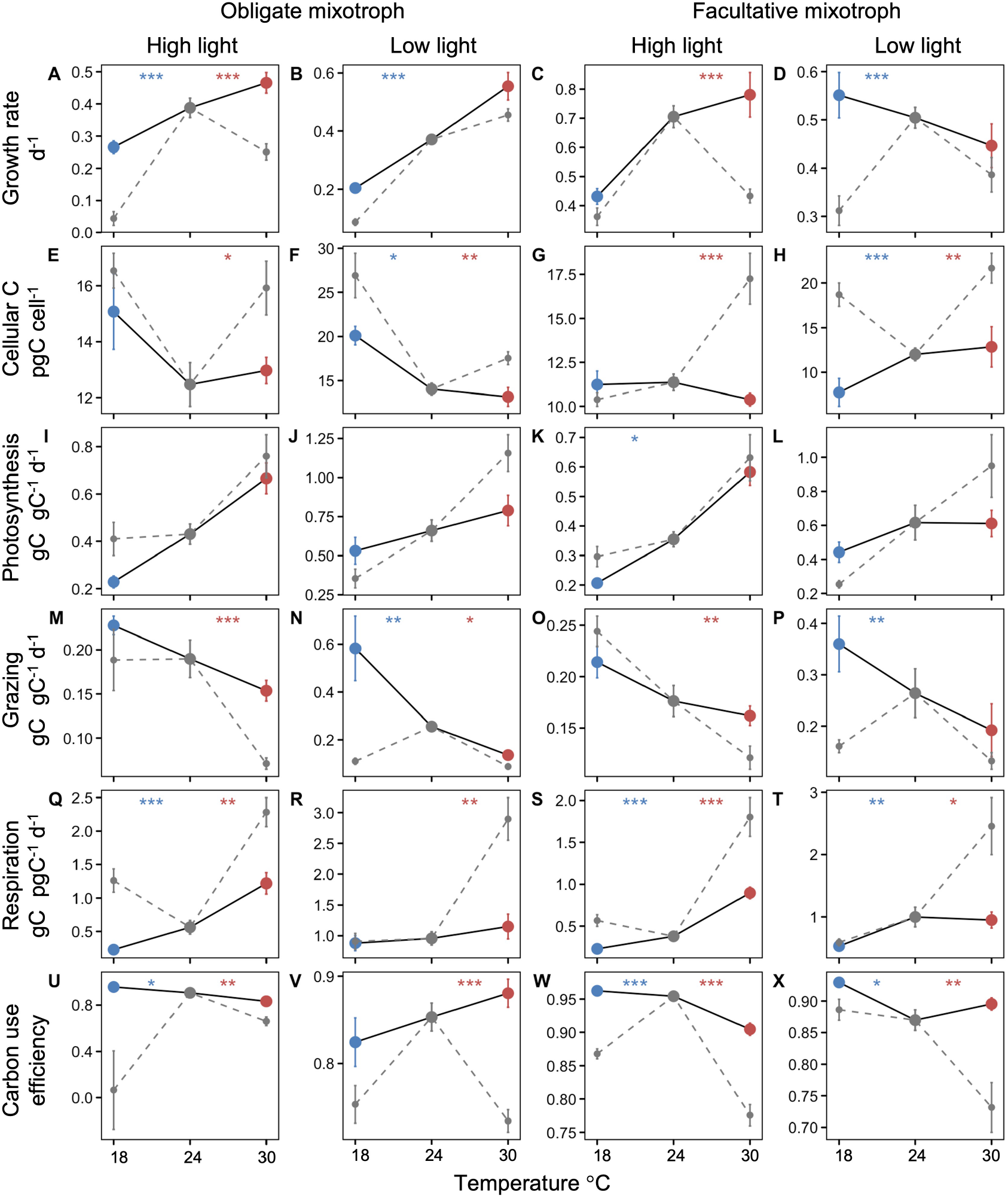
Thermal responses of carbon cycle-relevant traits in mixotrophs. We measured the plastic response of control lineages acclimated to all temperatures (gray points, dotted lines) and the evolved response of experimental lineages at their treatment temperatures (cold-evolved in blue, hot-evolved in red, solid black line). Data for the obligate (Strain 1393, first and second column) and the facultative mixotroph (Strain 2951, third and fourth column) are shown for both light levels (high light in first and third column, low light in second and fourth column). Points represent means for all 6 replicates across the final year of the project (2-3 time points) of the experiment, with error bars showing ± 1 standard error. We measured growth rate (A-D), cellular carbon content (E-H), photosynthesis (I-L), grazing (M-P), respiration (Q-T), and carbon use efficiency (U-X). Significant differences between the evolved and acclimated response at treatment temperatures are shown at the top of each panel in blue for 18°C and red for 30°C (two-sample Student’s t-test; *** = *p*<0.001, ** = *p*<0.01, and * = *p*<0.05).

Evolutionary responses in photosynthesis, grazing, and respiration were more variable. The thermal reaction norms of photosynthesis were most affected by evolution at the lowest light level, becoming less steep (higher photosynthetic rates at cold temperatures; lower photosynthetic rates at hot temperatures) after three years of evolution (Fig. 3, row 3). This may have resulted in part from a similar flattening of thermal reactions norms of chlorophyll at low light levels (Fig. S3, panels B and D), and due to changes in the use of photosynthetic machinery, especially increases in photosynthetic efficiency at low temperatures (Fig. S3). Grazing rates decreased slightly with temperature in evolved lineages, and thermal reaction norms were flatter in high light treatments (Fig. 3, row 4). Respiration rates decreased in evolved strains compared to acclimated control strains (Fig. 3, row 5). Collectively, this resulted in marked increases in carbon use efficiency across evolved lineages (Fig. 3, row 6).

### 3.3 Mechanisms underlying adaptation in obligate mixotroph at high light

For the obligate mixotroph at high light (which had the strongest evidence for adaptive evolution), metabolic and transcriptomic differences indicate evolutionary changes to cellular processes that may underlie adaptation. In the hot-evolved obligate mixotroph, increased growth rates at the hot temperature (Fig. 4A) were driven by a 19% reduction in cell size (Fig. 4B), a 116% increase in carbon intake from grazing (Fig. 4E), and a 46% decrease in respiratory costs (Fig. 4F) relative to the acclimated control. Smaller cell sizes mean more cells can be made with less carbon, which came from increased grazing and decreased respiratory loses. Although chlorophyll content normalized to cell size increased in the hot-evolved lineages (Fig. 4C), total carbon fixation was slightly lower (Fig. 4D). In cold-evolved lineages, cell sizes and grazing rates did not vary significantly from the control at cold temperatures (Fig. 4B and E), but increased growth rates may have been driven by an 80% decrease in respiratory costs (Fig. 4F).

**Figure 4.**
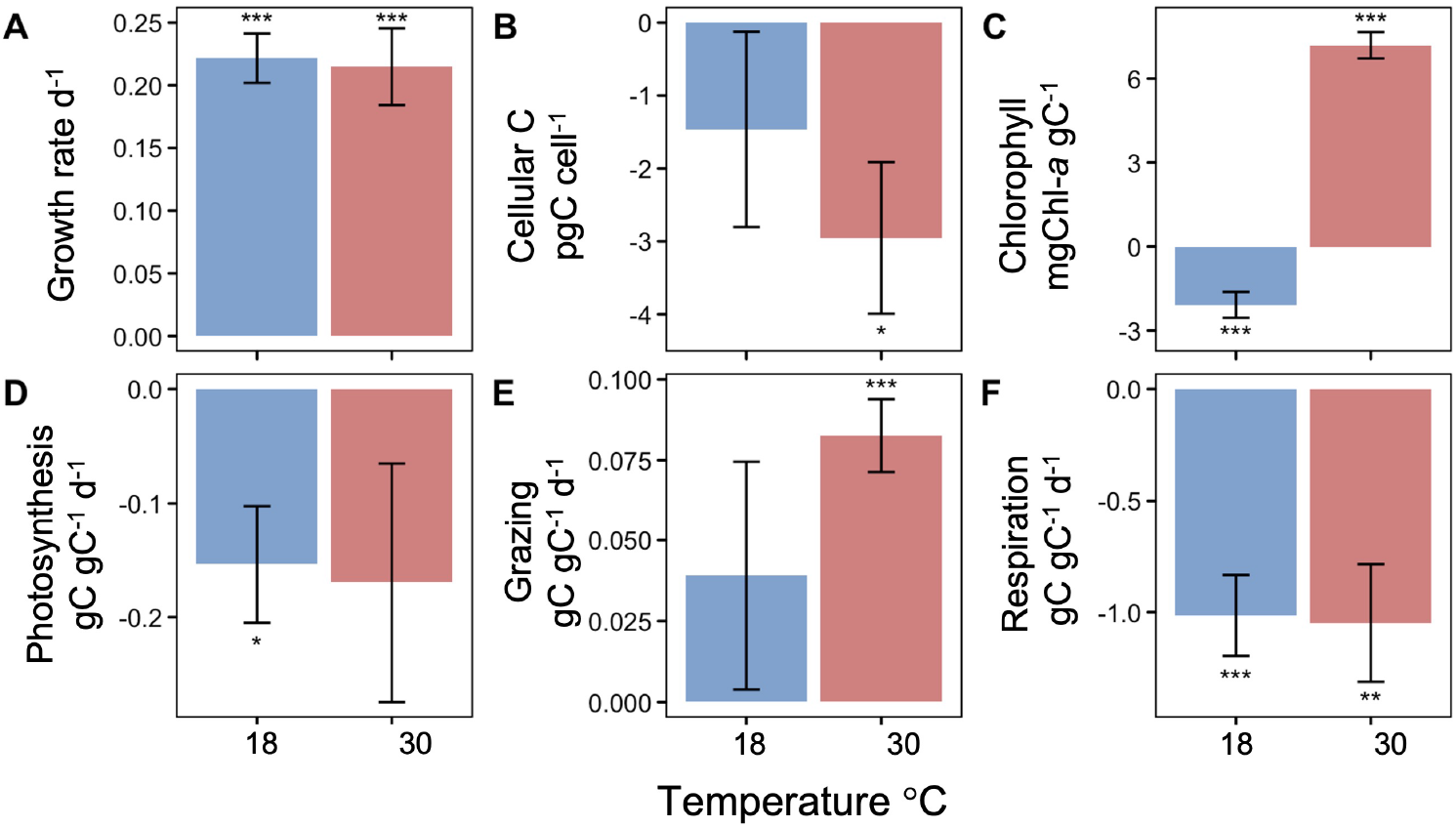
Differences between evolved and plastic responses of obligate mixotroph at high light. Differences between evolved responses of the cold and warm evolved lineages and the plastic responses of the acclimated controls at the respective cold and warm temperatures (obtained from reciprocal transplant) are shown for several key traits, to examine what mechanisms drove adaptation in the obligate mixotroph (Strain 1393) at high light (100 μmol quanta·m^−2^·s^−1^). The differences in cold-evolved lineages and acclimated controls are shown in blue, and the differences in hot-evolved lineages and acclimated controls is shown in red. Data are averaged for all 6 replicates across the last year of the experiment (2-3 time points) with error bars showing +/- 1 standard error. We measured growth rate (panel A), cellular C content (panel B), chlorophyll (panel C), photosynthesis (panel D), grazing (panel E), and respiration (panel F). Significant differences are shown by each bar (one-sample Student’s t-test; *** = *p*<0.001, ** = *p*<0.01, and * = *p*<0.05).

In the obligate mixotroph at high light, cold- and hot-evolved lineages showed changes in the thermal reaction norms for two important photosynthetic traits. Hot-evolved lineages had lower ETR per carbon at nearly all temperatures than those evolved at lower temperatures (Fig. 5A), but maintained photosynthetic efficiency to higher temperatures (Fig. 5B). The photosynthetic rates of cold-evolved lineages were more sensitive to changes in temperature (had steeper initial thermal response curve of ETR) and generally had lower photosynthetic efficiency (Fig. 5B). The grazing thermal response curves showed some signs of shifts in thermal optima but had much higher variability between replicates within the same treatment (Fig. S4).

**Figure 5.**
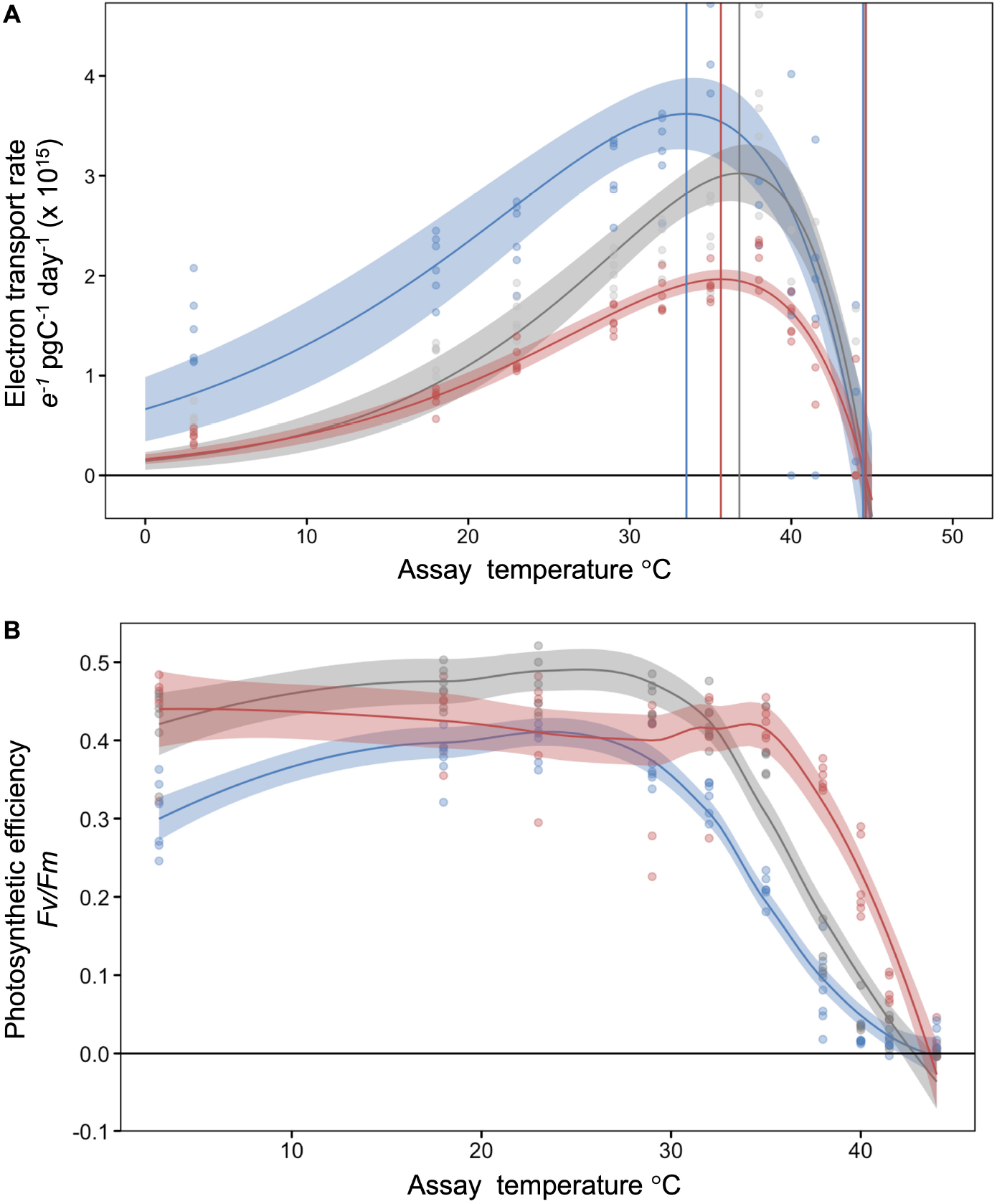
Thermal response curves of two photosynthetic traits for obligate mixotroph at high light. Electron transport rate per pgC (panel A) and photosynthetic efficiency (panel B) were measured at 10 temperatures between 3°C and 44°C for cold-evolved (blue), control (gray), and hot-evolved (red) lineages at the termination of the evolution experiment. For the cold-evolved lineages this represents >200 generations, and for the hot-evolved lineages, this represents >600 generations. Curves with confidence intervals represent the average of all 6 lineages at each temperature and individual lineages are represented by points. Three vertical lines farthest to the left represent the thermal optimum of each temperature, while the vertical lines on the right represent the thermal maxima.

Finally, we used our transcriptome data to understand changes in gene expression underlying our observed physiological responses. First, we confirmed that variation amongst our biological replicates exceeded any variation captured by technical replication (Fig. S5). We then proceeded with a comparative analysis using only one transcriptome per lineage. We found that, although some variation in gene expression existed across biological replicates, gene expression varied strongly by treatment (Fig. S6, Table S1). Of the 17,140 *Ochromonas* gene transcripts that we identified, we found that 6,951 genes were differentially expressed between lineages evolved at and adapted to 30°C. Of these, 380 genes had >2-fold upregulation, and 147 had >2-fold downregulation. Of 5,420 genes differentially expressed between lineages evolved at and adapted to 18°C, 105 had >2-fold upregulation, and 142 had >2-fold downregulation. Among these differentially expressed genes, we found evidence across metabolic pathways for a return to “baseline” gene expression levels over evolutionary time (Fig. 6). Specifically, cultures that experienced a 7-day acclimation to new temperatures tended to exhibit down-regulation when expression levels were compared to lineages evolved and acclimated at 24°C (Fig. 6). In contrast, evolved lineages had broadly similar gene expression levels across temperatures (Fig 6, S6).

**Figure 6.**
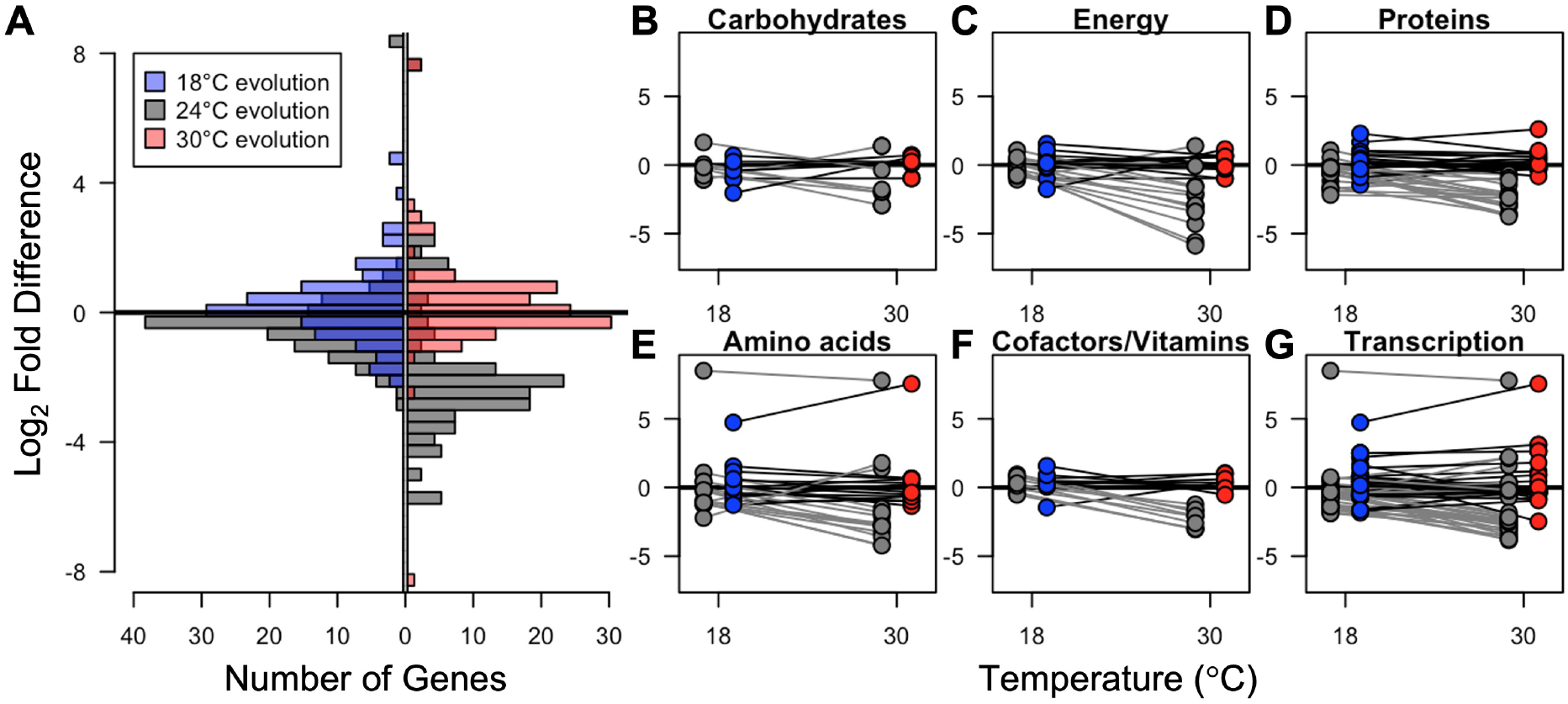
Evolution returns gene expression to homeostasis. **A:** Differences in expression of annotated genes between evolved and acclimated lineages at 18°C (left) and 30°C (right) and the control (24°C) lineages. Across genes that were significantly differentially expressed, we observed a tendency towards downregulation in acclimated lineages (gray; left side are acclimated to 18°C, right side are acclimated to 30°C), while evolved lineages tended to have gene expression levels similar to one another regardless of temperature (blue and red). **B-G:** This pattern was evident in many metabolic pathways (lines connect data from individual gene candidates), especially those associated with energy metabolism and biosynthesis. Colored points represent evolved lineages, and gray points represent control lineages acclimated to hot and cold temperatures.

## 4. Discussion

Mixotrophs play an integral role in oceanic food webs and the biological carbon pump (Mitra *et al*. 2014; Worden *et al*. 2015; Caron 2016; Ward & Follows 2016) and are predicted to become more heterotrophic with rising temperatures (Allen *et al*. 2005; Wilken *et al*. 2013). However, such predictions fail to account for evolutionary responses. We found that mixotrophs, like other unicellular organisms with fast generation times (Kawecki *et al*. 2012), can adapt to new thermal conditions within 50 generations, but that the magnitude of adaptation varied by mixotroph identity and environmental conditions. Generally, mixotrophs evolved lower respiration rates and higher carbon use efficiencies, responses that paralleled similar evolution experiments conducted in phytoplankton (Padfield *et al*. 2016; Schaum *et al*. 2018; Barton *et al*. 2020). At higher temperatures, mixotrophs evolved lower rates of photosynthesis and higher rates of grazing, compounding metabolic scaling predictions that mixotrophs will become more heterotrophic as temperatures increase (Rose & Caron 2007; Wilken *et al*. 2013; Princiotta *et al*. 2016).

Two competing processes shaped the consequences of mixotroph evolution on carbon cycling. On the one hand, evolved lineages had lower photosynthetic rates, higher grazing rates, and smaller cell sizes compared to control lineages at hot temperatures (Figs. 3, 4), suggesting mixotrophs could compound carbon dioxide atmospheric accumulation. On the other hand, evolved lineages also exhibited reduced respiration and higher carbon use efficiencies (Figs. 3, 4), suggesting that mixotrophs could increase trophic transfer efficiency and, potentially, carbon export (Ward & Follows 2016). However, we were unable to balance the mixotrophs’ carbon budget as carbon uptake (through photosynthesis and grazing) did not consistently match the sum of respiration and growth. One possible missing flux is the loss of organic carbon through leakage or exudation (Thornton 2014). Because we did not monitor pH evolution (except to confirm that pH did not change appreciably during exponential growth phase), alkalinity, or dissolved carbon within our cultures, we could not quantify this loss or how much carbon dioxide was absorbed through diffusion. Evidence suggests that *Ochromonas* likely do not have a carbon concentrating mechanism (CCM, Maberly *et al*. 2009), although transcriptomic analysis shows Strain 1393 retains the genes coding for proteins related to a CCM (Lie *et al*. 2018).

In general, evolved metabolic rates shifted back towards ancestral rates as they adapted, suggesting a recovery from stress-induced dysregulation to homeostasis. Stress has been shown to cause physiological and metabolic dysregulation in marine organisms (Fernández-Pinos *et al*. 2017, Innis *et al*. 2021). In our experiment, when mixotrophs were briefly acclimated to new thermal environments, they exhibited similar dysregulation of metabolism (either increases or decreases in metabolic rates relative to the 24°C control lineages; Fig. 3, gray lines) and gene expression (reduced expression; Fig. 6A, gray histograms). Yet over evolutionary time, mixotrophs adapted to the altered temperatures, such that evolved thermal reaction norms were flatter than acclimatized ones (Figure 3, compare “flatter” gray to “steeper” black lines) and relative gene expression levels returned to the optimized expression of the control (Figure 6A, blue and red histograms). This suggests that mixotrophs experience short-term acclimations as a “shock” that induces a stress response, but over time evolution allows them to recover by returning to an adaptive steady state homeostasis. The return to homeostasis in transcription regulation parallels other microbial systems (Brauer *et al*. 2008; López-Maury *et al*. 2008), including thermal response in *Escherichia coli* (Ying *et al*. 2015), suggesting global transcriptome optimization is a key component of adaptive thermal evolution.

Although all the mixotrophic lineages we evolved exhibited some evolutionary responses (Fig. 3), only the obligate mixotroph (*Ochromonas* Strain 1393) showed consistent evidence for adaptation (Fig. 2). The facultative mixotroph’s (Strain 2951) higher innate phenotypic plasticity may have resulted in a more muted evolutionary response: If the temperatures tested in our evolution experiment fell within Strain 2951’s capacity for plastic responses, this strain could have experienced reduced selection pressure compared to Strain 1393 (Snell-Rood *et al*. 2010). These findings intersect with a larger literature exploring the relationship between phenotypic plasticity and rapid evolution: More plastic lineages may be better able to survive in changing environments, thus “buying time” to evolve new adaptations (West-Eberhard 2003). But plasticity may also inhibit evolution when it inhibits the fixation of adaptive traits (Whitlock 1996). Because our study used only two mixotroph strains, we urge caution in interpreting our results in this context. However, constitutive mixotrophs fall along a wide spectrum of phenotypic plasticity, so future work could use this system as a testbed for these ideas.

Mixotroph evolution may also have been constrained by selection pressures imposed by our experimental design. For example, our experiment was conducted using warm water-adapted species (from an ancestral temperature of 24°C) that may have already been near the upper limits of thermal tolerance (Thomas *et al*. 2012). Thus, while the cold-evolved lineages of the obligate mixotroph shifted their photosynthetic thermal optima to slightly lower temperatures, the thermal optima for hot-evolved lineages did not change, and the thermal maxima were similar for all evolved lineages (Fig. 5). This supports the idea that thermal maxima that are physiologically constrained by metabolic limits are strongly phylogenetically conserved (Araújo *et al*. 2013). We also conducted our experiment in replete nutrient conditions and under stable temperature and light environments. In reality, nutrient limitation—which is likely experienced by mixotrophs in oligotrophic gyre habitats—can inhibit evolutionary adaptation (Thomas *et al*. 2017; Marañón *et al*. 2018; Aranguren-Gassis *et al*. 2019), as can the combination of multiple stressors (Brennan & Collins 2015). Additionally, our evolving lineages were xenic. Thus, bacterial prey in our experiment coevolved with the *Ochromonas* lineages. While this “community evolution” is a realistic scenario in marine ecosystems, this means that evolutionary changes in bacterial prey could have created complex feedbacks in prey availability and palatability.

In sum, our findings highlight the complex interaction between mixotroph identity and environmental selection pressures in constraining marine plankton adaptation. While some general trends (increased carbon use efficiency; flattening of thermal reaction norms; return to homeostatic gene expression) emerged, evolutionary responses were highly context dependent. These results suggest that incorporating evolutionary responses of marine microbes into climate predictions will be challenging. Additional studies may allow us to better link organismal metabolic plasticity to evolutionary responses and develop a more robust framework to predict the structure and function of upper ocean communities.

## Acknowledgements

We thank Sandeep Venkataram for discussions that shaped study design. We gratefully acknowledge manuscript feedback from Suzana Leles, Erika Eliason, Debora Iglesias-Rodriguez, and the members of the Moeller Lab at UC Santa Barbara. We thank Ryan Marczak, Gina Barbaglia, and Veronica Hsu for laboratory assistance, and the members of the Eliason Lab at UC Santa Barbara for lending us equipment and helping with early methods development. This work was funded by the National Science Foundation under grant NSF-BEE 185119 (to HVM).

## Author Contributions

MLB and HVM designed the study, conducted experiments, analyzed data, and wrote the manuscript. CP conducted transcriptomic lab work and analyses. CMM and EE developed protocols and conducted experiments. All authors contributed to manuscript revisions. The authors declare no conflict of interest.

## Supplemental Figures

**Figure S1.**
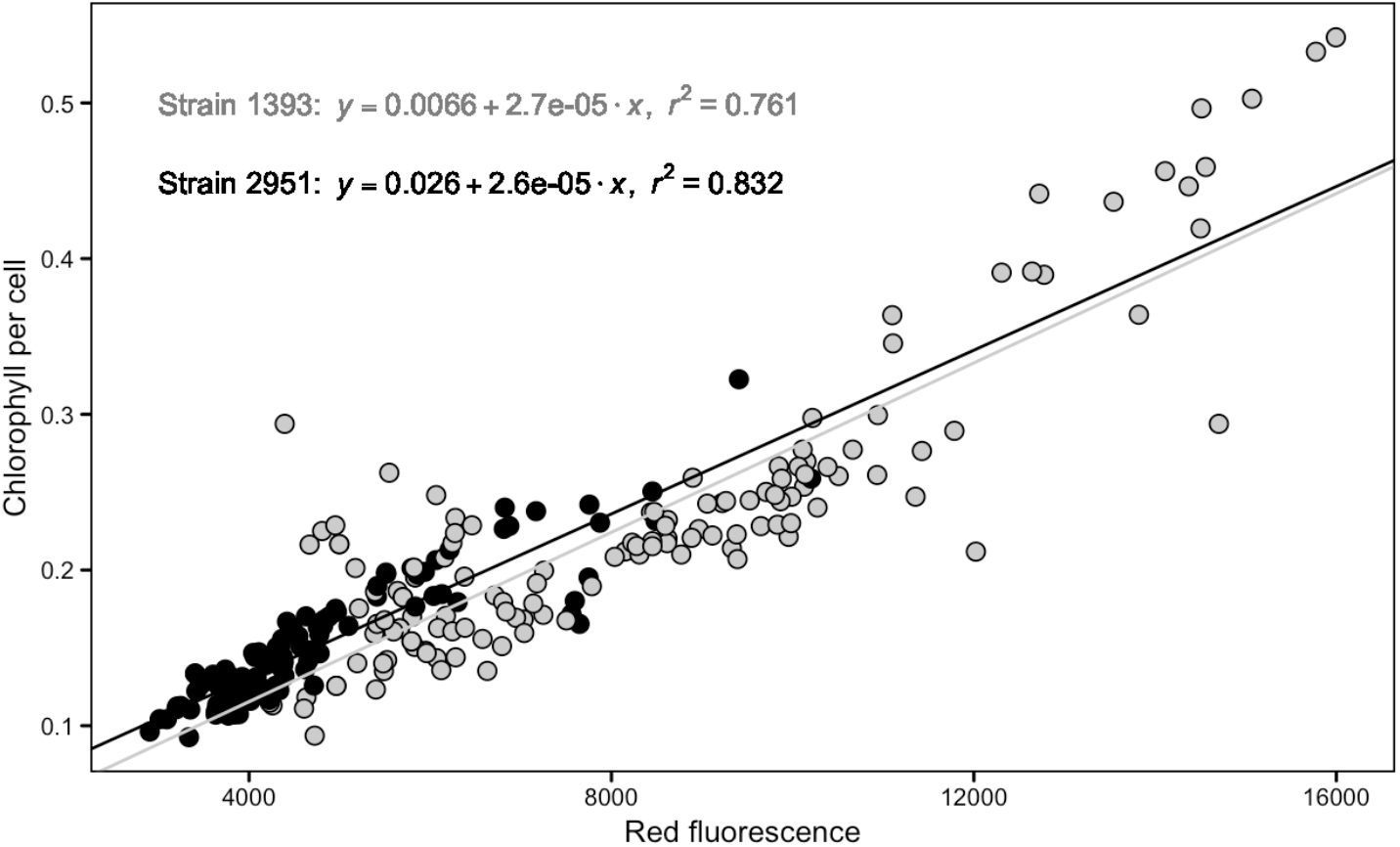
Correlation between extracted chlorophyll per cell measurements from a fluorometer and red fluorescence from flow cytometer for each strain of *Ochromonas* (obligate mixotroph Strain 1393 in gray, and facultative mixotroph Strain 2951 in black).

**Figure S2.**
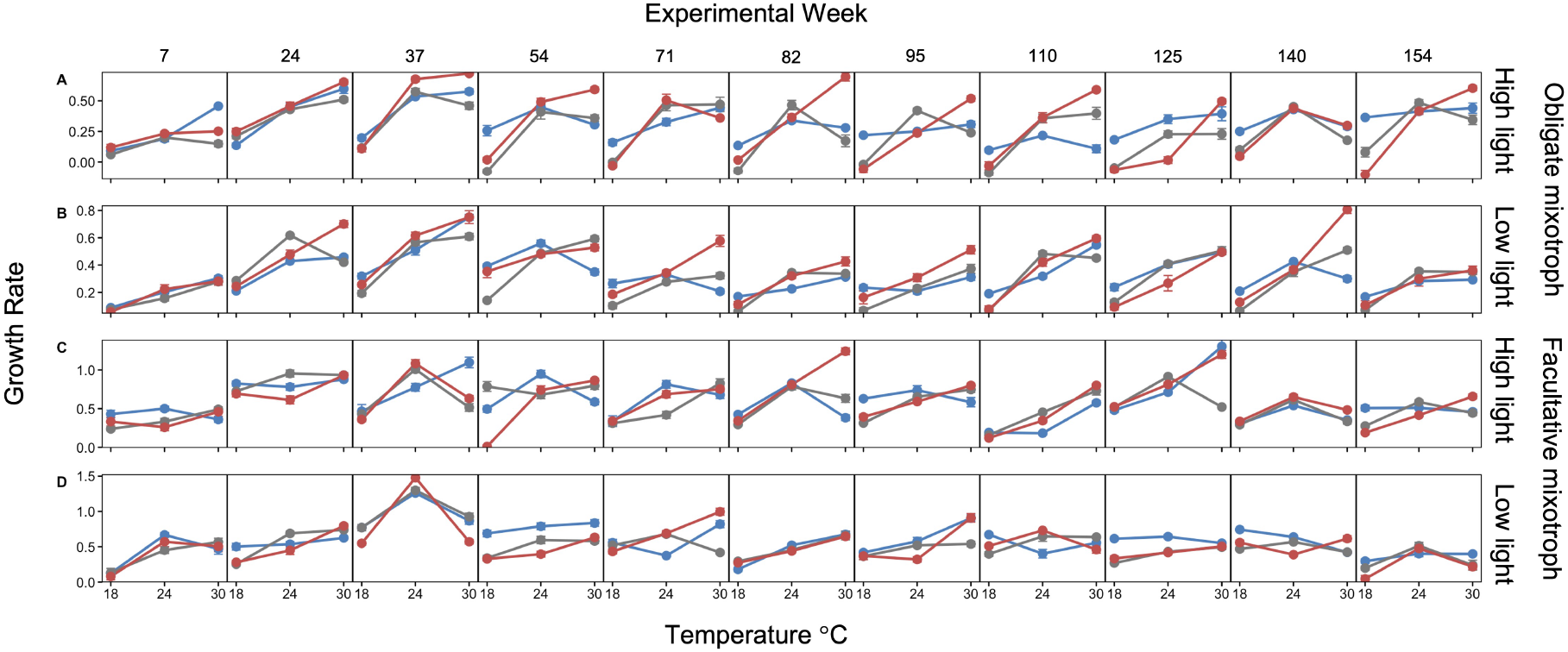
Thermal reaction norms of growth rate for all lineages at every reciprocal transplant time point. Panel A shows the obligate mixotroph at high light and panel B at low light. Panel C is the facultative mixotroph at high light and panel D at low light. Each box is a single time point labeled as experimental week going from left to right, with the x-axis of each box showing temperature. Blue points and lines are cold-evolved linages, red points and lines are hot-evolved lineages, and gray points and lines are control lineages. All points are averages of 6 replicates with bars depicting standard error.

**Figure S3.**
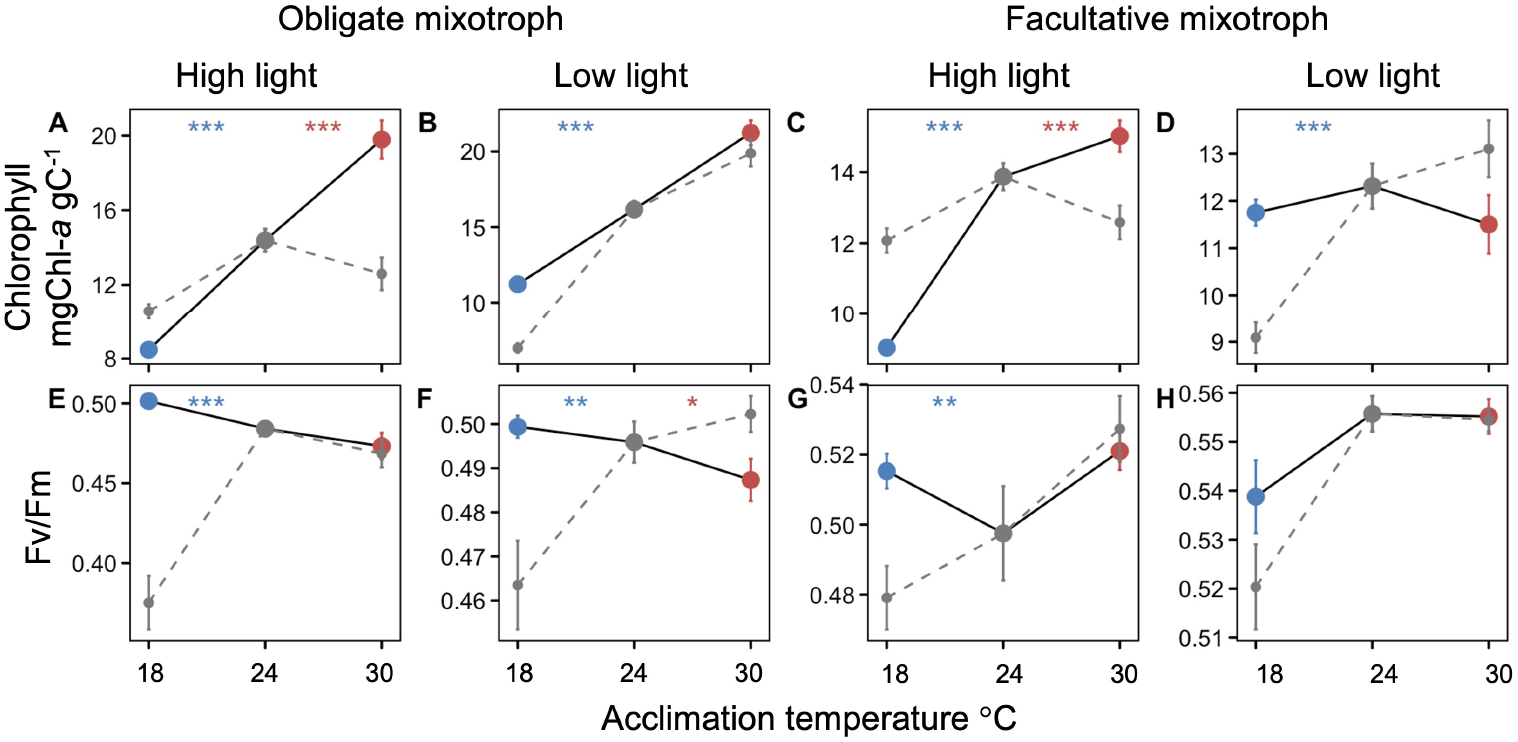
Additional thermal reaction norms of mixotroph traits before and after exposure to temperature treatments for evolutionary time. Control thermal reaction norms are in gray with dotted lines and post-adaptation data are shown as cold in blue and hot in red, connected by solid black lines. Top row has chlorophyll per pgC, and bottom row has photosynthetic efficiency (Fv/Fm). Points represent means for all 6 replicates across the final year of the project (2-3 time points) of the experiment, with error bars showing ± 1 standard error. Significant differences between the evolved and acclimated response at treatment temperatures are shown at the top of each panel in blue for 18°C and red for 30°C (two-sample Student’s t-test; *** = *p*<0.001, ** = *p*<0.01, and * = *p*<0.05).

**Figure S4.**
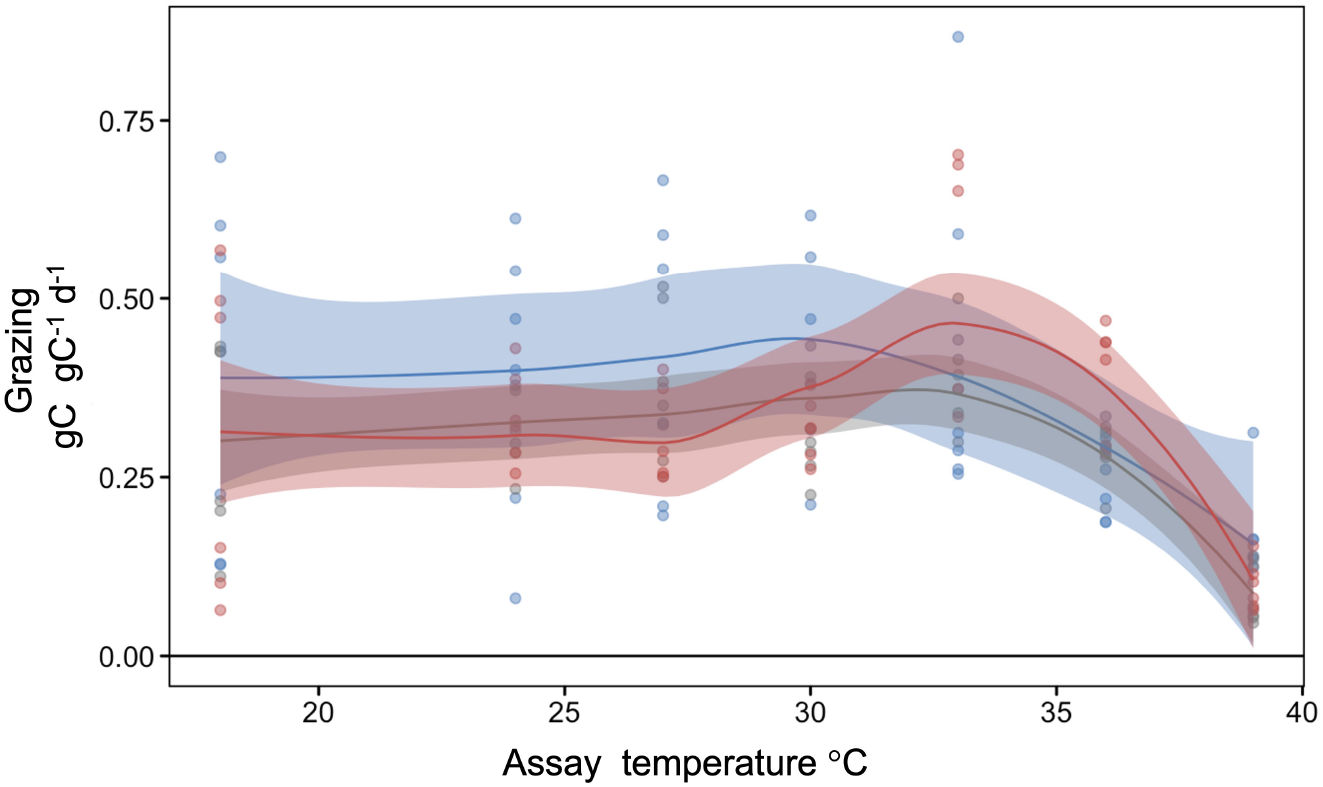
Thermal response curves of grazing rates for obligate mixotroph at high light. Grazing rate was measured at 10 temperatures between 18°C and 39°C for cold-evolved (blue), control (gray), and hot-evolved (red) lineages at the termination of the evolution experiment. For the cold-evolved lineages this represents >200 generations, and for the hot-evolved lineages, this represents >600 generations. Curves with confidence intervals represent the average of all 6 lineages at each temperature and individual lineages are represented by points. Three vertical lines farthest to the left represent the thermal optimum of each temperature, while the vertical lines on the right represent the thermal maxima.

**Figure S5.**
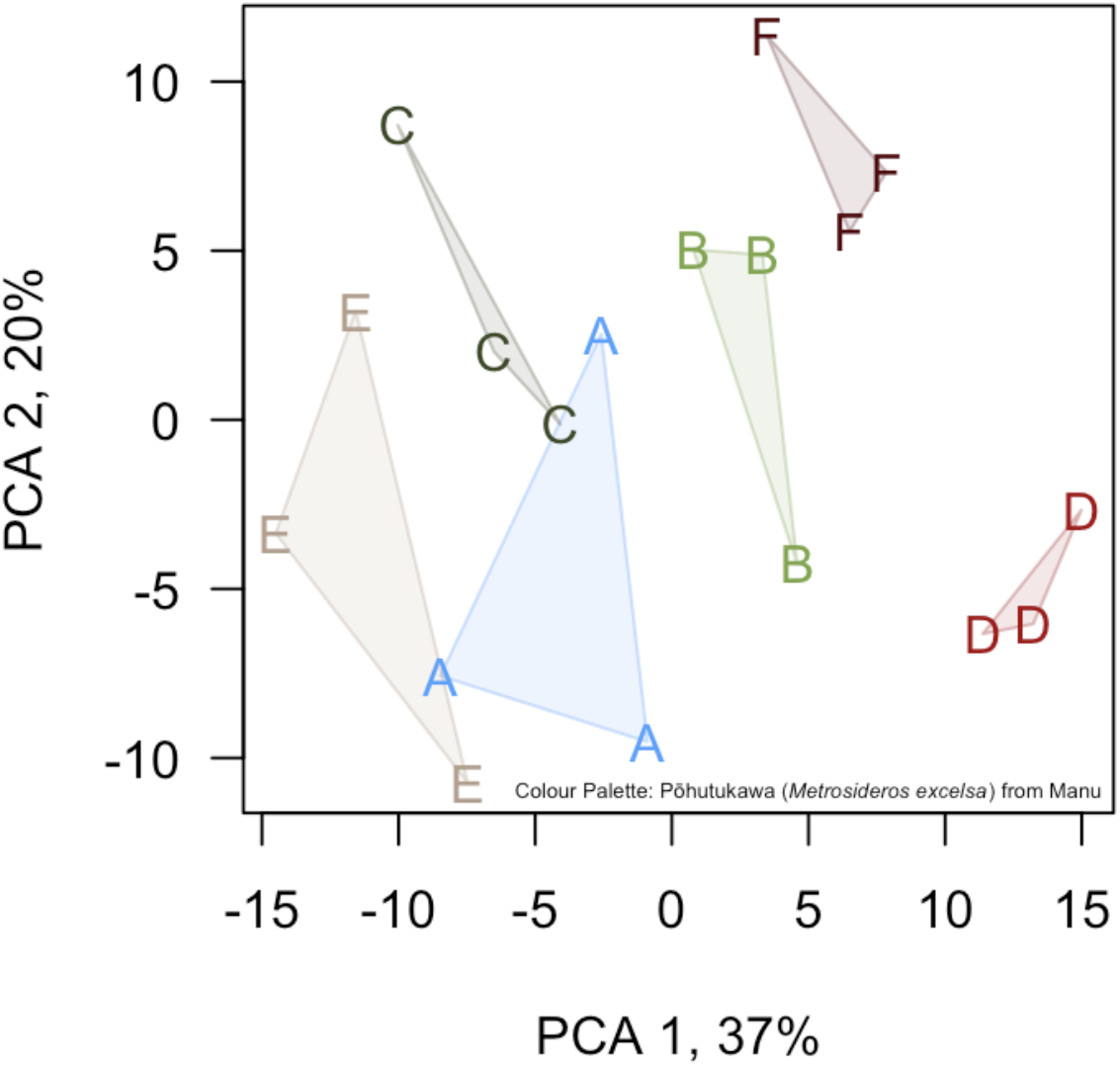
Biological replicates display more variation in gene expression than technical replicates. Data are drawn from *Ochromonas* lineages evolved at 18°C. Three technical replicates are shown for each of six biological replicates (labeled A through F).

**Figure S6.**
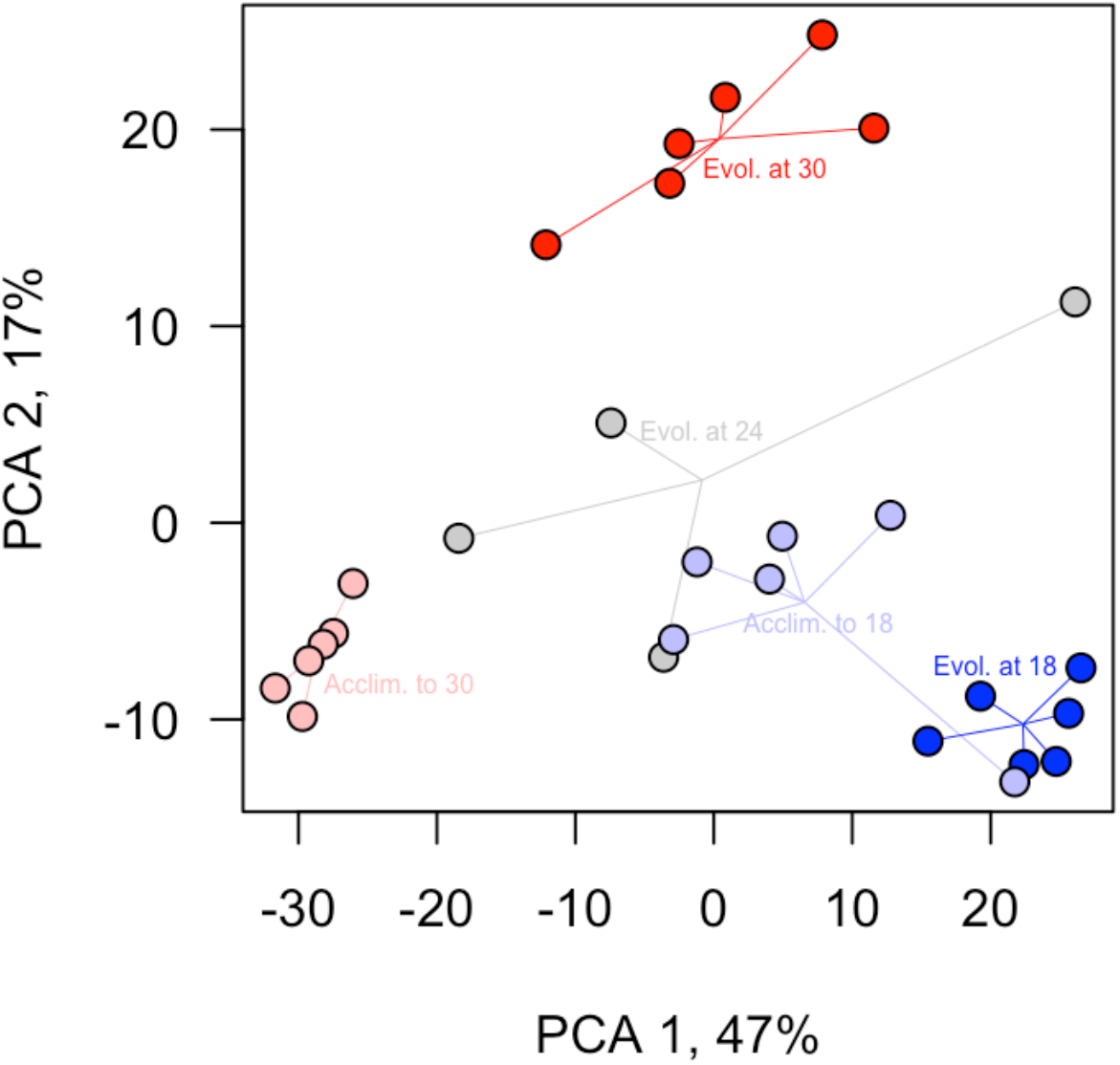
Principal component analysis of the 500 genes with the largest variance after variance stabilizing transformation, showing a strong treatment effect with replicates by treatment group. Generally, gene expression in *Ochromonas* Strain 1393 lineages evolved at 100 μmol quanta·m^−2^·s^−1^ reflected thermal environments. While control lineages evolving at 24°C and then acclimated to 18°C (light blue) showed intermediate gene expression between the control (gray; evolving at 24°C) and cold-evolved (dark blue) lineages, lineages evolved at (red) or acclimated to (light red) 30°C had distinctive expression patterns. Treatments are color coded with lines to the group mean.

**Figure S7.**
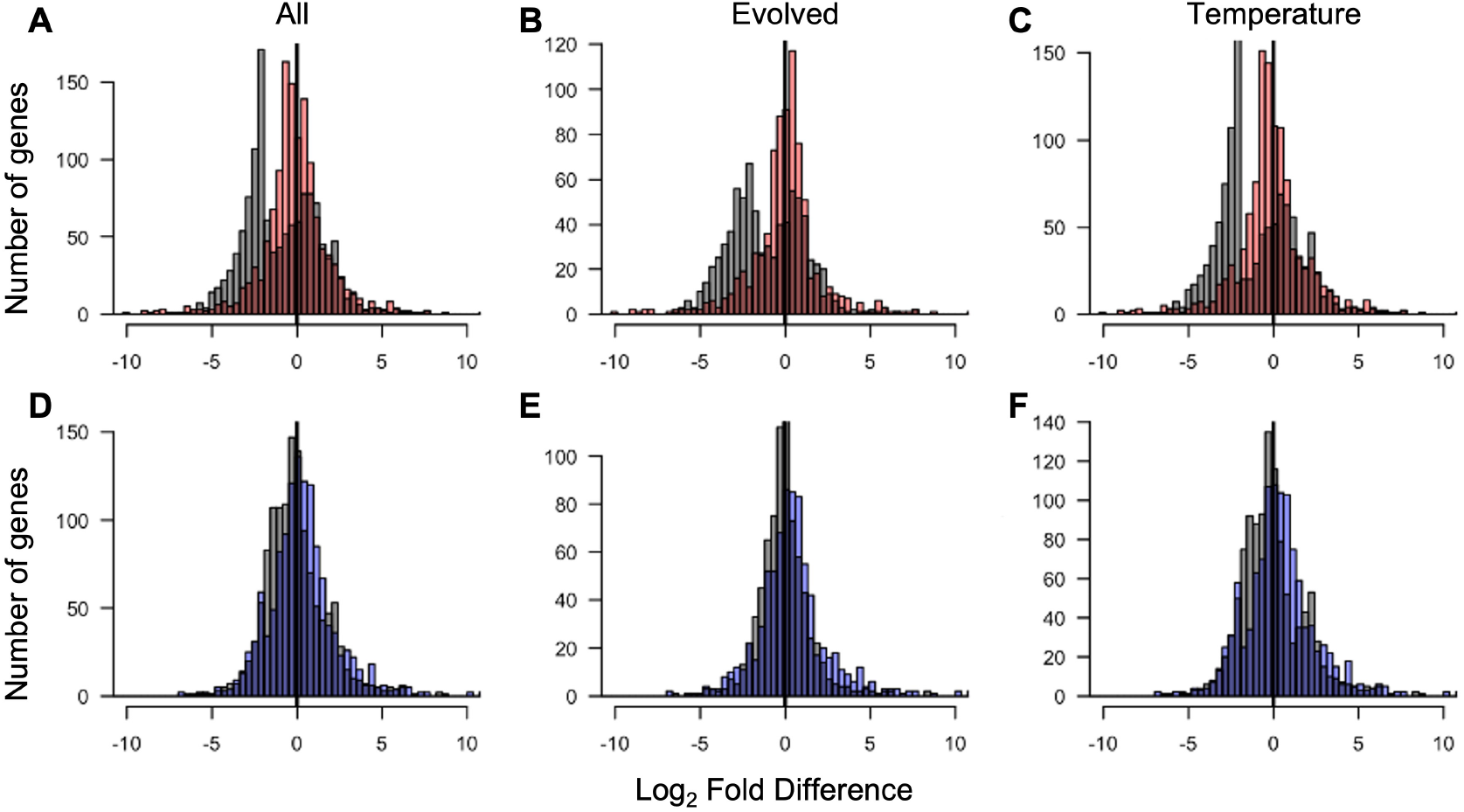
Regardless of gene subset, acclimated lineages display dysregulation while evolved lineages exhibit a return to homeostasis. Histograms are shown for all genes with significant differential expression; gray = acclimated lineages, red = evolved at 30°C, blue = evolved at 18°C. Left column: all differentially expressed genes. Middle column: Genes differentially expressed between lineages evolved or acclimated to 30°C or 18°C. Right column: Genes differentially expressed between lineages evolved and acclimated at 24°C and those evolved or acclimated at any other temperature. Top row: data from strains evolved at or acclimated to 30°C. Bottom row: data from strains evolved at or acclimated to 18°C.

**Table S1.** Number of differentially expressed genes (two-fold difference with adjusted p-value < 0.1) in *Ochromonas* Strain 1393 lineages evolved at high light. Rows show differences in assay temperature, and cell entries show number of genes either up or down-regulated for a series of pairwise comparisons (columns). Here, evolved = lineages evolved and assayed at treatment temperature; control = lineages evolved and assayed at 24°C; and acclimated = lineages evolved at 24°C and assayed at 18°C or 30°C. Rows of pairwise comparisons sum to greater than the final column because some genes are differentially expressed across multiple pairwise comparisons.

|  | Evolved compared<br>to control |  | Acclimated<br>compared to control |  | Evolved compared<br>to acclimated |  | All unique<br>differential genes |  |
| --- | --- | --- | --- | --- | --- | --- | --- | --- |
| Temperature | up | down | up | down | up | down | up | down |
| 18°C | 121 | 141 | 100 | 124 | 124 | 118 | 284 | 257 |
| 30°C | 95 | 84 | 147 | 380 | 88 | 482 | 498 | 644 |

## Notes

https://github.com/mleporibui/OchEvo

### Competing Interest Statement

The authors have declared no competing interest.

